# Modelling belowground plant acclimation to low soil nitrogen – An eco-evolutionary approach

**DOI:** 10.1101/2025.05.16.653651

**Authors:** Arjun Chakrawal, Sacha J. Mooney, Tino Colombi

## Abstract

Increased root growth to access greater soil mineral nitrogen resources and increased root exudation to stimulate microbial mineralisation of soil organic nitrogen are widely observed plant acclimations to nitrogen limitation. However, their quantitative contribution to plant growth and ecosystem productivity remains largely elusive. Here, we present a novel optimality-based eco-evolutionary model in which plants dynamically regulate carbon partitioning between root growth and exudation to maximise their aboveground growth. Our simulations indicated that the dynamic availability of soil mineral and organic nitrogen as well as plant nitrogen demand and nitrogen uptake capacity shape optimal carbon partitioning between root growth and exudation. The simulated carbon allocation patterns aligned with empirical studies on belowground plant responses to varying soil nitrogen resources. Our eco-evolutionary approach represents a paradigmatic change in modelling plant nitrogen foraging, which is essential to generate hypotheses on optimal plant acclimation in future soil environments characterised by more erratic nitrogen availability.

## 1. INTRODUCTION

Nitrogen (N) limitation is a major constraint to the primary productivity of terrestrial ecosystems (Du *et al*. 2020; LeBauer & Treseder 2008; Liu *et al*. 2010). As an indispensable component of chlorophyll, N is essential to photosynthesis and thus carbon (C) assimilation by plants. On the other hand, N foraging by plants requires C translocation from above-to belowground plant tissues. Hence, the coupled C and N fluxes in plant-soil systems are key drivers for plant growth and primary productivity of terrestrial ecosystems (Körner 2015). It is expected future soil environments will be characterised by more erratic N availability. Increased frequency and severity of dry and wet spells will lead to more irregular, less predictable microbial N mineralisation and denitrification patterns and accelerate leaching of mineral N (Ollivier *et al*. 2011; Rennenberg *et al*. 2009). Moreover, tightening environmental regulations limiting the use of mineral fertiliser and rising fertiliser prices will likely decrease the abundance of readily plant-available mineral N in agricultural soils (Liu *et al*. 2010; Lynch *et al*. 2022).

Except for nodule-forming plant species that live in symbiosis with nitrogen-fixing bacteria, soil-derived, primarily mineral N is the only N resource directly available to plants. Root-colonising mycorrhizal fungi can significantly contribute to plant N uptake (Courty *et al*. 2015), yet around 15% of plant species do not form mycorrhizal symbiosis (Genre *et al*. 2020). Almost all vascular plants develop a (fine) root system to directly access and take up mineral N from soil (Nacry *et al*. 2013). Furthermore, through the exudation of organic compounds, plants can stimulate microbial mineralisation of organic N in the vicinity of their roots to gain access to soil organic N resources (Cheng *et al*. 2014; Phillips *et al*. 2011). To acclimate to low soil N availability, plants increase C supply to their root system to promote root growth (Gedroc *et al*. 1996; Lopez *et al*. 2023; Nacry *et al*. 2013) or increase C exudation from roots into the surrounding soil (Darwent *et al*. 2003; Phillips *et al*. 2011). The projected trend towards more erratic N availability in soil (Liu *et al*. 2010; Lynch *et al*. 2022; Ollivier *et al*. 2011; Rennenberg *et al*. 2009) suggests these acclimation mechanisms will be key for plant growth in future soil environments.

The stimulation of root growth and root exudation requires greater C translocation from the aboveground to the root system, which increases the C costs of soil exploration (Brunn *et al*. 2022; Darwent *et al*. 2003; Hodge 2006). In turn, greater root growth and exudation enable plants to access larger soil N resources and to take up more N from soil. This close coupling of C and N fluxes shows plants must increase the amount of N taken up per C invested into root growth and exudation to maximise their growth under N limiting conditions. Therefore, the ratio between C investment into root growth and exudation and the resulting N uptake determines the acclimative value of these belowground responses for plant growth and ecosystem productivity (Colombi *et al*. 2024; Lynch 2015). However, quantification of C and N fluxes in plant-soil systems and underlying physiological processes that characterise the N foraging behaviour of plants is challenging and laborious, especially under field conditions (Cusack *et al*. 2024; Jones *et al*. 2009; McCormack *et al*. 2015). Therefore, quantitative data on the effects of increased root growth and exudation on plant growth under N limiting conditions remain scarce. Mathematical models can —at least partially— help overcome these challenges (Ahkami *et al*. 2024; Wang *et al*. 2023), thereby providing novel insights into the role of belowground plant acclimations for plant growth and ecosystem productivity.

Several models that simulate effects of different N foraging strategies on plant growth and productivity have been developed. Structural-functional plant growth models link root architecture and anatomy with carbon, nutrient, and water fluxes at the single plant scale (Postma *et al*. 2017; Schnepf *et al*. 2018). Other models such as the Fixation and Uptake of Nitrogen (FUN) model simulate the effects of different N uptake pathways on coupled C and N fluxes in plant-soil systems at the field and ecosystem scale (Brzostek *et al*. 2014; Fisher *et al*. 2010; Sulman *et al*. 2017). However, these models are all based on pre-determined root phenotypes or prescribed C allocation patterns to different N acquisition pathways (e.g. root growth and exudation). Therefore, they do not allow to explore how plant growth can be maximised through acclimations in N foraging behaviour. Optimality-based eco-evolutionary approaches, which assume that plants traits are optimised in a way that enables maximum productivity under given environmental conditions (Harrison *et al*. 2021; Martiny *et al*. 2023), may overcome this reliance on fixed root physiological behaviour. For example, eco-evolutionary modelling has been used to show how optimal ligninolytic activity maximises microbial growth during plant litter decomposition (Chakrawal *et al*. 2024). Modelling N foraging with an eco-evolutionary approach allows us to elucidate how plants maximise their growth by regulating C partitioning between root growth and exudation. Thus, eco-evolutionary modelling can uncover N foraging strategies underpinning efficient plant acclimation to more erratic soil N availability.

Here, we present a novel parsimonious N foraging model that explicitly accounts for the ability of plants to acclimate to varying soil N resources. In our model, plant N acquisition is formulated as an optimal control problem where plants regulate C partitioning between root growth and exudation to maximise aboveground biomass over a given growth period. We then applied this eco-evolutionary approach to elucidate how the soil’s capacity to provide N and the demand and ability of plants to take up N affects C allocation in plant-soil systems.

## 2. MATERIAL AND METHODS

### 2.1. Structure of nitrogen foraging model

We employed a minimalistic parsimonious modelling approach for simulating plant growth coupled with microbially mediated soil organic matter (SOM) decomposition during vegetative development of annual plants. For simplicity, we assumed that plants acquire N directly from the soil solution and biological N fixation and N uptake via mycorrhizal fungi were excluded. Aboveground plant tissue, roots biomass, root exudates, soil microorganisms, SOM, and soil mineral N were represented as single pools (Figure 1A). Furthermore, we assumed that belowground C investment and N mineralisation and uptake happen in the uppermost 30 cm of soil. Bulk density was set to 1.2 g cm^-3^, which is typical for topsoil across land uses (Panagos *et al*. 2024). All C and N fluxes were expressed as daily rates on a per-area basis, i.e. g C m^-2^ d^-1^ and g N m^-2^ d^-1^. Mass balance equations coupling C and N dynamics and kinetic relationships between C stocks and rates are provided in Supplemental Table S1 and S2, respectively.

**Figure 1.**
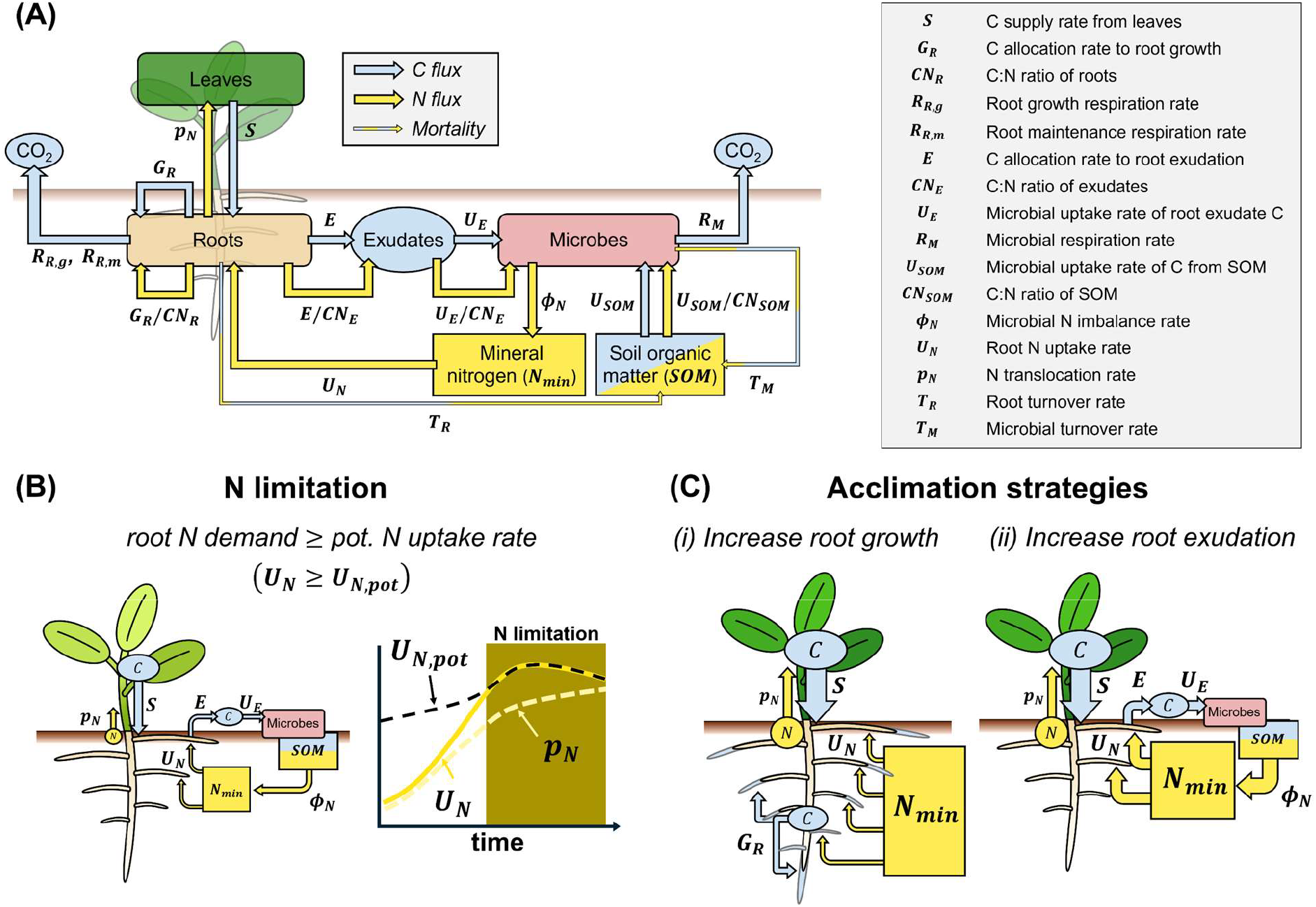
Schematic representation of (A) nitrogen foraging model with key carbon and nitrogen fluxes and pools in plant-soil systems, (B) definition of nitrogen limitation at the root level occurring when root nitrogen demand (*U*_*N*_) exceeds potential root N uptake rate (*U*_*N*,*pot*_), and (C) belowground plant acclimation to nitrogen limitation through (i) increasing root growth and (ii) increasing root exudation.

Using big leaf approximation (Bonan *et al*. 2021), aboveground plant tissues were represented as a single leaf. We assumed gross photosynthesis (*A*) to be nitrogen limited and we used a logistic function to simulate gross photosynthesis as a function of leaf N stock with additional effect of internal shading (Ågren 1985). Leaf respiration (*R*_*L*_) and net C assimilation of the plant (*A*_*net*_) were calculated using a fixed leaf carbon use efficiency (*CUE*_*L*_). The portion of *A*_*net*_ allocated belowground was defined as the root carbon supply (*S*). A fraction of *S* was allocated to root growth respiration (*R*_*R*,*g*_), as determined by the root carbon use efficiency (*CUE*_*R*_). The remaining proportions of *S* was partitioned among root growth (*G*_*R*_), maintenance respiration (*R*_*R*,*m*_), and root exudation (*E*). We used equilibrium chemistry approximation kinetics (Tang & Riley 2013) to describe microbial uptake of exudated C (*U*_*E*_) and SOM (*U*_*SOM*_). The partitioning of microbial C uptake into growth and respiration was described by a fixed microbial carbon use efficiency (*CUE*_*M*_). Plant and microbial necromass was assumed to recycle into the SOM pool at rates *T*_*R*_, and *T*_*M*_, respectively (Figure 1A).

Carbon and nitrogen fluxes were coupled using C:N ratios of the different pools. For soil microorganisms, we assumed a plastic C:N ratio to reflect their ability to increase their C:N ratio upon N limitation (Manzoni *et al*. 2021). Nitrogen imbalance fluxes (*ϕ*_*N*_) resulting from differences in C:N ratio between soil microorganisms, SOM, and root exudates indicated net microbial N mineralisation (*ϕ*_*N*_ > 0) or immobilisation (*ϕ*_*N*_ < 0). We assumed C:N homeostasis for plants (Körner 2015; Nacry *et al*. 2013), represented by fixed C:N ratios for leaves, roots, and root exudates. Moreover, we assumed that plants take up only mineral N (*U*_*N*_) from which a part is translocated aboveground to meet leaf N demand (*p*_*N*_), while the remaining N is used for root growth and exudation (Figure 1A).

### 2.2. Definition of N limitation

We defined N limitation at the root level. Therefore, N limitation occurs when root N demand (*U*_*N*_) exceeds potential root N uptake (*U*_*N*,*pot*_) due to low availability of mineral N in the soil (*U*_*N*_ ≥ *U*_*N*,*pot*_; Figure 1B). We defined *U*_*N*,*pot*_ as a function of available mineral N and root N using a Michaelis-Menten kinetics as,

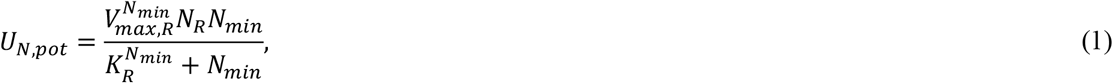

where *N*_*min*_ and *N*_*R*_ are the soil mineral N and root N stocks, respectively, and 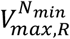 and 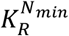 are the maximum rate constant for mineral N uptake by roots and half saturation constant, respectively. *U*_*N*_ was defined as the sum of N demand for leaf growth, root growth and exudation, and can be written as,

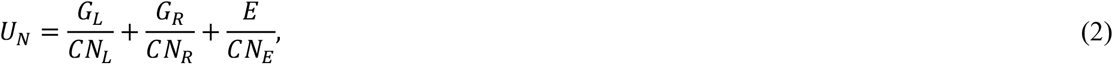

where, *G*_*L*_, *G*_*R*_, and *E* denote carbon allocation to leaf growth, root growth, and root exudation, respectively, and *CN*_*L*_, *CN*_*R*_, and *CN*_*E*_ denote C:N ratios of leaves, roots, and exudated, respectively. Note, the N demand for leaf growth equals the translocation of N from the roots to aboveground plant tissues, i.e., 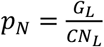.

We defined *G*_*L*_ and *G*_*R*_ as,

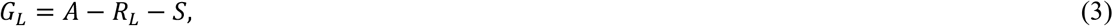

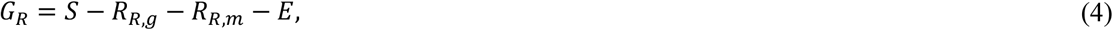

Detailed definitions of *R*_*L*_, *R*_*R*,*g*_ and *R*_*R*,*m*_ are provided in Supplemental Table S2. The eco-evolutionary approach used to estimate *E* and *S* is described in the following section.

Since N limitation was defined by the logical condition *U*_*N*_ ≥ *U*_*N*,*pot*_, root N uptake under N limiting conditions was set to potential root N uptake, i.e., *U*_*N*_ = *U*_*N*,*pot*_. To maintain C:N homeostasis of leaves under N limitation, plants must regulate leaf growth rate to match the reduced N uptake rate. In our model, plants can achieve this by increasing C supply to roots, which in turn leads to greater root growth and/or increased root exudation. Due to the root C:N homeostasis condition, root N (*N*_*R*_) increases upon greater root growth, which in turn increases *U*_*N*,*pot*_ (Eqn. 1). Increased root exudation enhances N mineralisation and thus soil mineral N (*N*_*min*_), leading to greater *U*_*N*,*pot*_ (Eqn. 1). Hence, we directly linked leaf and root C:N homeostasis conditions with enhanced root growth and exudation and thus key plant acclimation strategies to N limitation (Figure 1C)(Darwent *et al*. 2003; Gao *et al*. 2015; Gedroc *et al*. 1996; Henry *et al*. 2007; Krapp *et al*. 2011; Liljeroth *et al*. 1990; Lopez *et al*. 2023; Nacry *et al*. 2013; Phillips *et al*. 2011; Vives-Peris *et al*. 2020).

### 2.3 Eco-evolutionary approach to capture plant acclimation dynamics

Given the coupling of C and N fluxes in plant-soil systems, the regulation of C partitioning between root growth and root exudation is required to maximise cumulative aboveground plant growth. To explicitly account for this regulation, we formulated our nitrogen foraging model as an optimal control problem (OCP; see Supplemental Table S3 for full mathematical description). Thereby, the model finds the optimal C partitioning between root growth and exudation that maximises cumulative aboveground plant growth during a given growth period with C and N mass balance (Table S1) and plant C:N homeostasis as dynamic constraints. The objective function (*J*) was as follows,

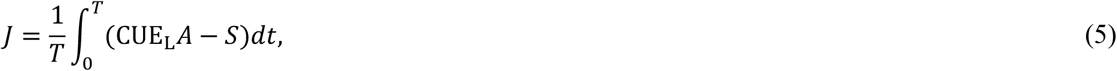

where, *T* is the growth period. The OCP setup utilises root C supply and exudation as control variables to maximise cumulative aboveground growth. As a result of the optimisation process, we obtained optimal trajectories of C and N pools, along with their fluxes, including root C supply and root C exudation. From these, the optimal root growth rate was calculated using Eqn. 4.

Since the OCP simultaneously solves the state and control variable, nonnegativity constraints of C and N stocks were added. The condition for N limitation described above was implemented imposing the constraint 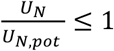, which ensured that root N uptake rate could not exceed the potential N uptake rate.\ To ensure physiologically and ecologically meaningful C partitioning patterns, we assumed i) that at least 20% of net assimilated C is allocated to leaf growth and to root C supply, respectively, and ii) that at least 20% of root C supply is allocated to root growth. Lastly, we imposed a maximum root exudation rate constraint, i.e., 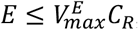, where 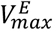 is the rate constant for maximum root exudation, and *C*_*R*_ is the root C. The OCP was solved numerically with the MATLAB toolbox ‘Yop’ that uses direct collocation method (Leek 2016).

### 2.4 Model parametrisation and initial conditions

Due to data availability, we parametrised the model to represent C and N dynamics in maize systems. A detailed list of model parameters, their assigned values, and the initial conditions of the state variables is provided in the Supplemental Tables S4 and S5. We ran the model for 120 days, which is representative for the vegetative growth period of maize.

The parameters *a* and *b* in the expression for gross photosynthesis rate, *A* (Supplemental Table S2), were adjusted to approximate the observed range of gross primary production rate from maize (Jiang *et al*. 2021; Suyker & Verma 2012). For the leaf N carrying capacity parameter, *K*_*N*,*L*_, we assumed a value of 75 g N m^-2^. The C:N ratio of leaves and roots were fixed at 21 and 50, respectively (Zhang *et al*. 2020). To simulate exudation of sugars and organic acids, we assumed a very high C:N ratio of 10 000 for root exudates. The initial value of C:N ratio for microbial growth was set at 10, which was allowed to vary depending on N imbalance (Zhang & Elser 2017). The C:N ratio of SOM was set to 10 (Tipping *et al*. 2016). We assumed 20 g kg^-1^ SOM content with 0.04% of that as active microbial biomass C (Malik *et al*. 2018). Leaf, root, and microbial carbon use efficiencies were set to 0.7, 0.6, and 0.3, respectively (Blagodatskaya *et al*. 2014; Hansen *et al*. 2009; Herrmann & Colombi 2019; Mo *et al*. 2021). Microbial and root turnover rate were set to 5 year^-1^ (Ryan *et al*. 2020) and 0.1 year^-1^ (Gill & Jackson 2000), respectively.

We assumed that nitrate is the only mineral N species taken up by roots. Nitrate uptake rate kinetics of maize roots were used to parametrise the maximum rate constant for root N uptake 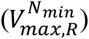 and half saturation constant for potential root N uptake 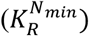. Based on the maximum root nitrogen uptake rate reported by York *et al*. (2016) of 110 μM NO ^-^ g^-1^ root dry weight h^-1^, we set root N content to 10 g N kg^-1^ root dry weight (Zhao *et al*. 2019), assumed 30 cm soil depth and 1.2 g cm^-3^ bulk density (Panagos *et al*. 2024), and set 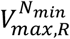 to 0.25 d^-1^. Furthermore, 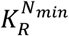 was assumed to be 20 g N m^-2^, which ensured that our simulated root nitrogen uptake rate kinetics aligned with empirical studies (Griffiths & York 2020; York *et al*. 2016).

The maximum root exudation rate constant 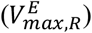 was set to 0.035 d^-1^, which ensured that the simulated root exudation fluxes (0.25 to 2 g C m^2^ d^-1^) were within the range of root exudate production fluxes reported in empirical studies (Chari *et al*. 2024; Phillips *et al*. 2008). To parametrise microbial uptake kinetics of exudates, we approximated exudates as carbohydrates (i.e. sugars) and organic acids. We selected kinetic parameters that ensured that the simulated exudate uptake rates (0.4 to 4 g C m^2^ d^-1^) aligned with observed microbial uptake rates of glucose (Rousk *et al*. 2014). Similarly, the parametrisation of microbial N uptake and SOM decomposition kinetics ensured that simulated microbial N uptake and SOM decomposition rates aligned with empirical data (Abramoff *et al*. 2022; Weintraub-Leff *et al*. 2023).

### 2.5. Simulating effects of soil and plant properties on optimal nitrogen foraging behaviour

To elucidate the capability of our model to depict effects of soil fertility on C and N fluxes underpinning optimal plant acclimation to N limitation, we simulated different capacities of the soil to provide N to plants. We combined SOM C:N ratios between 5 and 30 with different initial soil mineral N stocks (*N*_*min*_; 10 and 20 g N m^-2^) and slow and fast microbial SOM decomposition rates (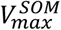 0.15 and 0.30 d^-1^). Differences in plant N demand were simulated through leaf C:N ratios of 15 and 30, representing high and low N demand. Finally, we used two different maximum rate constants for root N uptake to simulate plants with low and high N uptake capacity (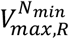 0.25 d^-1^ and 0.75 d^-1^).

## 3. RESULTS

### 3.1. Soil nitrogen resources modulate carbon allocation dynamics

We simulated three N availability scenarios to elucidate C allocation dynamics underpinning plant acclimation to different soil N resources: First, a base scenario with a high initial soil mineral N content (20 g N m^-2^) and a low SOM C:N ratio of 10 indicating a high potential for mineralisation of soil organic N; Second, greater plant reliance on soil mineral N due to a high initial mineral N content (20 g N m^-2^) and a limited potential for mineralisation of soil organic N (SOM C:N ratio = 20); And third, greater plant reliance on microbial mineralisation of organic N resources due to a low initial mineral N content (10 g N m^-2^) and a high potential for mineralisation of soil organic N (SOM C:N ratio = 10).

Differences in soil N resources altered C partitioning patterns throughout the simulated 120-day growth period. Compared to the base scenario, low mineral N content and limited N mineralisation potential decreased the fraction of net assimilated C allocated to leaf growth within the first week (Figure 2A), while root C supply increased (Figure 2C). The fraction of net assimilated C supplied to roots remained comparatively high throughout most of the growth period under low mineral N. In contrast, root C supply under limited N mineralisation potential converged with the base scenario as plant development progressed (Figure 2C). Upon completion of the growth period, the three scenarios showed distinctly different cumulative above-belowground C partitioning patterns. Compared to the base scenario, the ratio between C allocated to leaf growth and C supplied to roots decreased under limited N mineralisation potential and low mineral N. Thus, a shift towards greater belowground C allocation was needed to maximise cumulative aboveground growth under decreased soil N availability (Figure 2B and D).

**Figure 2.**
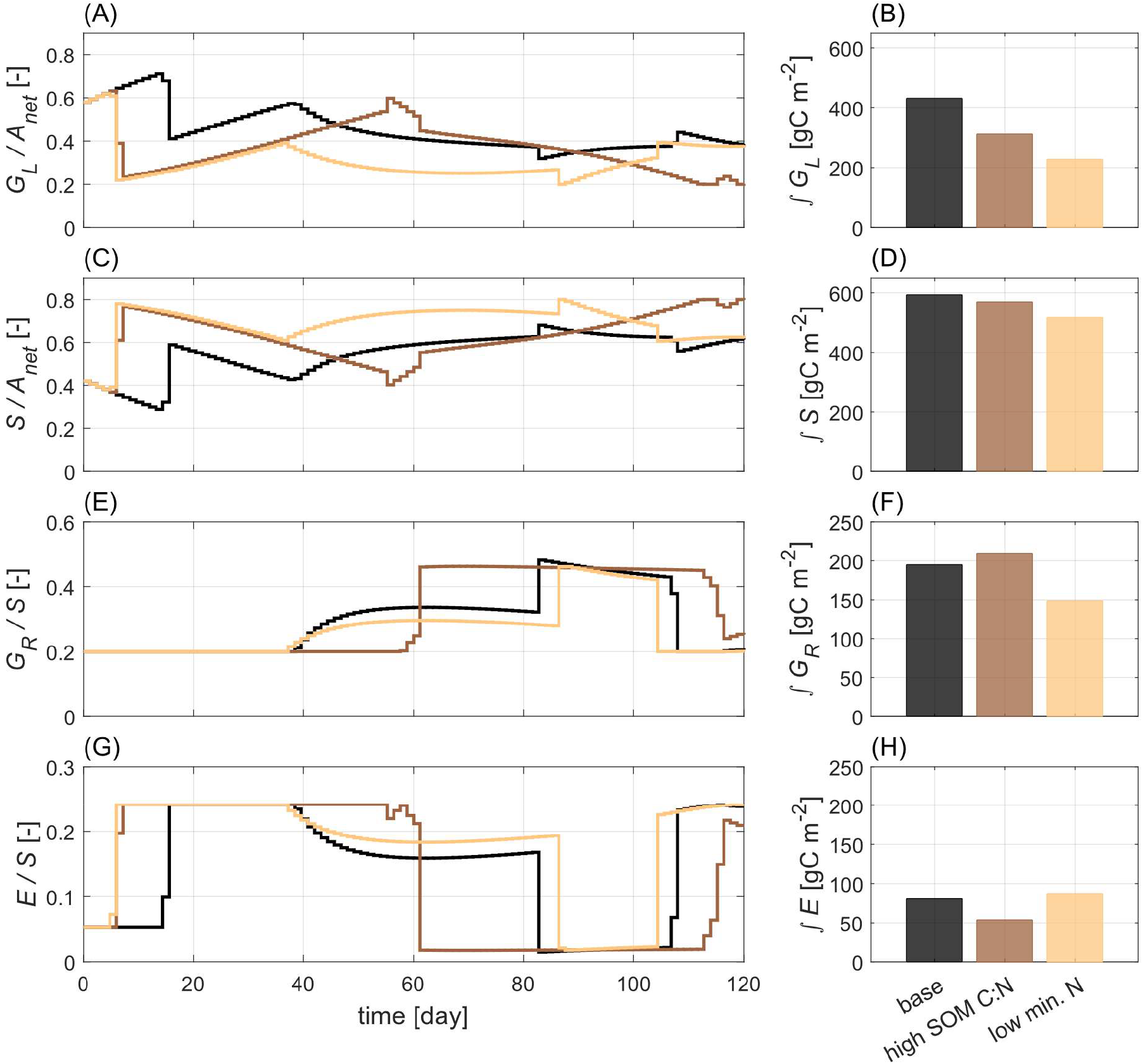
Effects of soil N resources on carbon allocaiton dynamics enabling maximum cummulative aboveground plant growth over 120-day growth period. (A) and (C) show daily fraction of net assimilated C (*A*_*net*_) alloacted to leaf growth (*G*_*L*_) and supplied to roots (*S*), respectively. (E) and (G) show daily fraction of *S* alloacted to root growth (*G*_*R*_) and root exudation (*E*), respectively. (B), (D), (F), and (H) show cummulative amount of carbon allocated to *G*_*L*_, *S, G*_*R*_, and *E* over 120-day growth period. Black line and bar denote high soil mineral N (20 g N m^-2^) and high potential for mineralisation of soil organic N due to a low soil organic matter carbon-nitrogen ratio (SOM C:N = 10); Brown line and bar denote high soil mineral N (20 g N m^-2^) and low potential for mineralisation of soil organic N due to a higher SOM C:N ratio (SOM C:N = 20); Beige line and bar denote lower soil mineral N (10 g N m^-2^) and high potential for mineralisation of soil organic N due to a low SOM C:N ratio (SOM C:N = 10).

Belowground C partitioning dynamics also showed distinct responses to alterations in soil N availability. Greater plant reliance on microbial mineralisation of organic N resources in soil required a comparatively high C allocation to root exudation throughout most of the simulated growth period. Under limited N mineralisation potential, however, a shift from C allocation to exudation towards root growth occurred after around half of the simulated growth period (Figure 2E and G). The cumulative C partitioning patterns resulting from these differences in belowground C allocation dynamics reflected differential plant acclimation strategies to varying soil N availability. The increased reliance on microbial mineralisation of organic N resources due to low soil mineral N required greater C allocation to root exudation to maximise cumulative aboveground plant growth. In contrast, greater reliance on mineral N resources due to a limited N mineralisation potential increased the importance of root growth for maximising cumulative aboveground growth (Figure 2F and H).

### 3.2. Low soil organic nitrogen mineralisation potential favours root growth

In addition to SOM quality, represented by SOM C:N ratio, SOM decomposition rates are key to the mineralisation of soil organic N. We included two different rate constants for microbial SOM uptake (slow: 0.15 d^-1^, fast: 0.30 d^-1^) to simulate this effect of microbial physiology on plant acclimation to different soil N resources over a 120-day growth period. Under low and high initial mineral N, faster SOM decomposition reduced the fraction of net assimilated C supplied to roots (Figure 3A and C). Belowground C partitioning patterns underlying maximum cumulative aboveground plant growth differed distinctly between SOM decomposition rates. With slow SOM decomposition, C allocation to root growth increased with SOM C:N ratio while C allocation to exudation decreased, especially under high soil mineral N content. Under fast SOM decomposition, however, such pronounced changes in belowground C allocation upon decreasing SOM quality did not occur. Carbon partitioning between root growth and exudation was only marginally affected by SOM C:N ratio under fast SOM decomposition, suggesting high soil microbial activity can at least partially compensate for poor SOM quality (Figure 3B and D).

**Figure 3.**
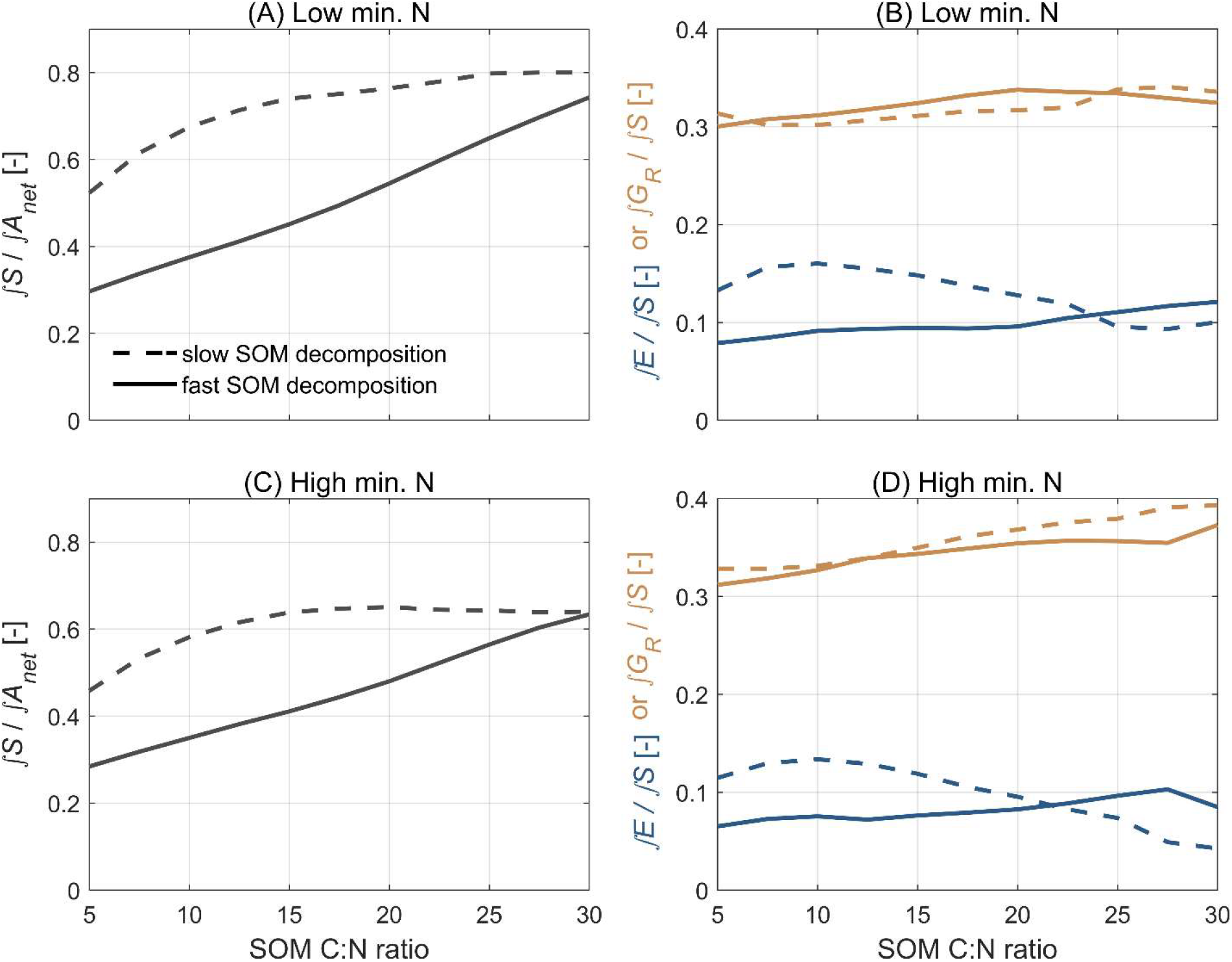
Effects of microbial soil organic matter (SOM) decomposition rate, soil mineral N, and SOM C:N ratio on C partitioning patterns in plant-soil systems underpinning maximum cumulative aboveground plant growth over simulated 120-day growth period. (A) and (C) show fraction of net assimilated C (*A*_*net*_) supplied to roots (*S*); (B) and (D) show fraction of *S* allocated to root exudation (*E*; blue lines) and root growth (*G*_*R*_; orange lines). Slow and fast microbial SOM decomposition rate refer to maximum rate constants for microbial SOM uptake of 0.15 and 0.30 d^-1^; Low and high soil mineral N refer to mineral N stocks of 10 and 20 g N m^-2^.

### 3.3. Interplay between nitrogen supply and demand shape belowground carbon partitioning

Variation of leaf C:N ratio allowed simulating the effects of plant N demand on optimal plant acclimation strategies maximising cumulative aboveground plant growth during the simulated 120-day growth period. High and low plant N demand were indicated by leaf C:N ratios of 15 and 30, respectively. The fraction of net assimilated C supplied to roots increased with plant N demand across different SOM C:N ratios and mineral N contents (Figure 4A and C). In contrast, belowground C partitioning patterns did not show consistent responses. Under low mineral N content and low plant N demand, a shift in C allocation from root exudation to root growth occurred upon increasing SOM C:N ratio. Similar, yet less pronounced patterns occurred under high mineral N content for high and low plant N demand (Figure 4B and D). Conversely, under low soil mineral N and high plant N demand, the importance of root exudation increased with SOM C:N ratio (Figure 4B). These distinct differences in simulated belowground C partitioning patterns pointed out the importance of the interplay between plant N demand and soil N supply for plant acclimation to different soil N resources.

**Figure 4.**
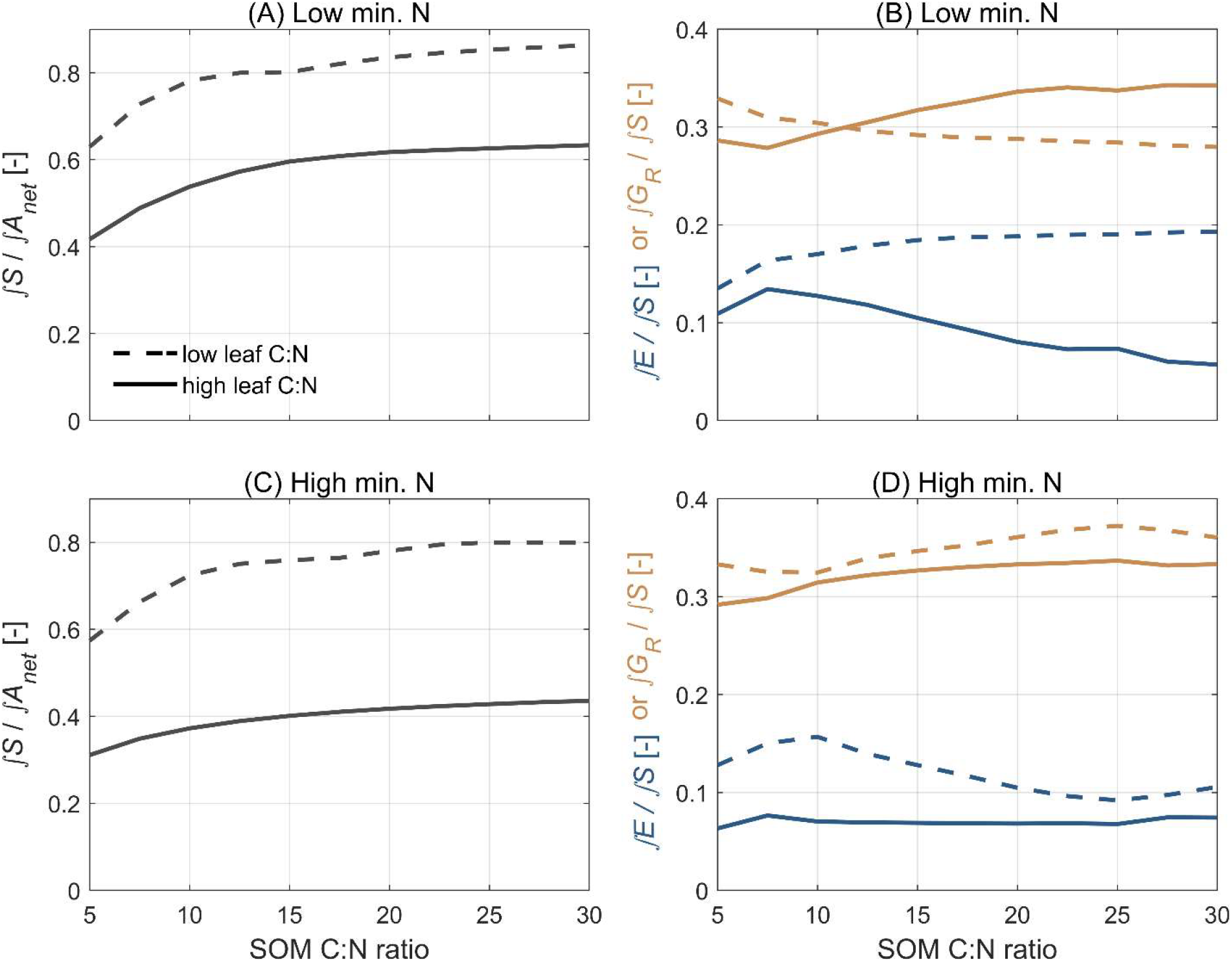
Effects of plant N demand, soil mineral nitrogen (N), and soil organic matter (SOM) C:N ratio on C partitioning patterns in plant-soil systems underpinning maximum cumulative aboveground plant growth over simulated 120-day growth period. (A) and (C) show fraction of net assimilated C (*A*_*net*_) supplied to roots (*S*); (B) and (D) show fraction of *S* allocated to root exudation (*E*; blue lines) and root growth (*G*_*R*_; orange lines). High and lows plant N demand refer to leaf C:N ratios of 15 and 30; Low and high soil mineral N refer to mineral N stocks of 10 and 20 g N m^-2^.

### 3.4. Importance of root exudation increases with root nitrogen uptake potential

To simulate the role of root N uptake capacity in plant acclimation to different soil N resources, we included two different maximum rate constants for root N uptake (low: 0.25 d^-1^; high: 0.75 d^-1^). With greater root N uptake capacity, a lower fraction of net assimilated C had to be supplied to roots to maximise cumulative aboveground plant growth over 120 days, regardless of soil mineral N content (Figure 5A and C). Belowground C partitioning patterns were also clearly affected by root N uptake capacity. The importance of root exudation for maximising cumulative aboveground plant growth increased with root N uptake capacity. This effect was particularly pronounced if the capacity of the soil to supply N was comparatively high due to a high mineral N content or a low SOM C:N ratio (Figure 5B and D).

**Figure 5.**
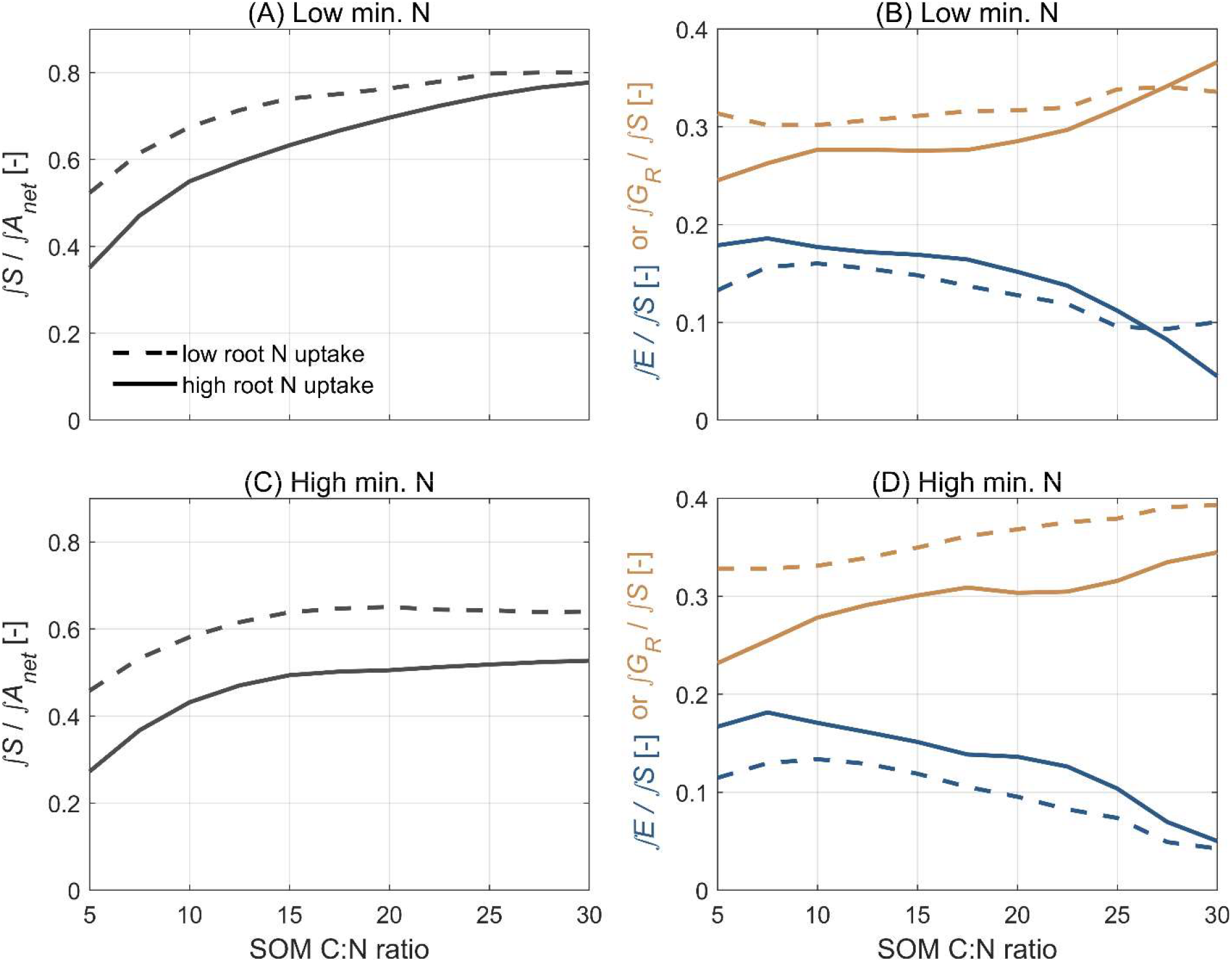
Effects of root N uptake rate, soil mineral nitrogen (N), and soil organic matter (SOM) C:N ratio on C partitioning patterns in plant-soil systems underpinning maximum cumulative aboveground plant growth over simulated 120-day growth period. (A) and (C) show fraction of net assimilated C (*A*_*net*_) supplied to roots (*S*); (B) and (D) show fraction of *S* allocated to root exudation (*E*; blue lines) and root growth (*G*_*R*_; orange lines). Low and high root N uptake rate refer to maximum rate constants for root N uptake of 0.25 d^-1^ and 0.75 d^-1^; Low and high soil mineral N refer to mineral N stocks of 10 and 20 g N m^-2^.

## 4. DISCUSSION

Here, we developed a novel parsimonious yet highly versatile N foraging model that explicitly acknowledges that ability of plants to acclimate their belowground physiology in response to variable soil N resources (Canarini *et al*. 2019; Colombi *et al*. 2024; Darwent *et al*. 2003). Adopting an eco-evolutionary approach allowed us to estimate optimal C partitioning between root growth and exudation as an outcome of the maximisation of cumulative aboveground growth. Thereby, our model does not rely on prescribed root physiological behaviour like other models simulating C and N fluxes in plant-soil systems (Brzostek *et al*. 2014; Fisher *et al*. 2010; Lokupitiya *et al*. 2009; Postma *et al*. 2017; Schnepf *et al*. 2018; Sulman *et al*. 2017). This capacity for dynamic regulation of C partitioning between root growth and exudation was the key to account for the pivotal importance of belowground plant acclimations to maximise N uptake and aboveground plant biomass.

Our simulations suggested that the maximisation of cumulative aboveground plant growth requires an early increase in root C supply (Figure 2A and C), which corresponds to decreasing root-shoot ratios during vegetative development observed in annual plants (Gedroc *et al*. 1996; Santangeli *et al*. 2024; Siddique *et al*. 1990). The early increase in root C supply coincided with greater C allocation to root exudation, reflecting the limited capacity of young, small root systems to access mineral N resources in soil. Over time, the relative contribution of root exudation for maximising cumulative aboveground growth decreased, while that of root growth increased, especially if the potential for mineralisation of organic N resources was limited (Figure 2E and G). These results agree with empirical studies reporting peak root exudation during early vegetative development (Aulakh *et al*. 2001; Chen *et al*. 2019). The simulated changes in the importance of root growth and exudation for N uptake and aboveground growth during plant development highlight that our model captures the dynamic nature of plant N foraging.

Increased root-shoot ratios is one of the most widely observed acclimation strategies to N limitation of non-leguminous plants (Gedroc *et al*. 1996; Lopez *et al*. 2023; Nacry *et al*. 2013). This was reflected in our simulations by the greater fraction of net assimilated C supplied towards roots in response to a reduced capacity of the soil to supply N, a greater plant N demand, and a lower N uptake capacity of plants (Figures 3-5, A and C). In roots, C partitioning patterns underpinning maximum cumulative aboveground growth were more nuanced. The combination of high SOM C:N ratio and slow SOM composition increased C allocation to root exudation. This corresponds to a recent metanalysis that reported decreased root carbon exudation in response to soil N addition (Zeng *et al*. 2024). However, if we performed the same simulations with fast SOM decomposition, C allocation to root exudation was almost unaffected by SOM C:N ratio, indicating that increased microbial activity can alleviate effects of poor SOM quality (Figure 3B and D). Under high plant N demand and low soil mineral N, representing a scenario of potentially severe N limitation, C allocation to root exudation even increased with SOM C:N ratio (Figure 4B). Previous studies showed that plants compensate limited N uptake capacity of roots with the development of a larger and denser root system (Dunbabin *et al*. 2004; York *et al*. 2016). Here, we found that C allocation to root growth required to maximise cumulative aboveground plant growth increased in response to lower N uptake capacity, which reflects this compensation (Figure 5B and D). These results demonstrate the capacity of our N foraging model to provide insights into the effects of soil and plant properties on the emergence of distinct root acclimation strategies.

In our model, we considerably simplified N acquisition by plants and only considered direct root N uptake. Hence, we did not account for plant N uptake resulting from symbiosis with rhizobia or mycorrhizal fungi, which are of significant importance for plant N acquisition (Courty *et al*. 2015). Similarly, we restricted our simulations to the vegetative development of annual plants and excluded effects of internal plant N remobilisation, which are particularly important during reproductive growth and in perennial plant species (Masclaux-Daubresse *et al*. 2010; Tegeder & Masclaux-Daubresse 2018). Since biological N fixation, mycorrhiza-mediated N uptake, and internal N remobilisation are fuelled by carbon fixed by plants through photosynthesis (Finlay 2008; Fisher *et al*. 2010; Tegeder & Masclaux-Daubresse 2018), the same optimisation principles apply to these processes as to root growth and exudation. In addition to root growth and exudation, plants must regulate C allocation to biological N fixation, mycorrhiza-mediated N uptake, and internal plant N remobilisation to maximise aboveground biomass. Therefore, our framework can readily be adapted to also include the effects of biological N fixation, mycorrhiza-mediated N uptake, and internal N remobilisation on plant growth under N limiting conditions.

Another simplification we made is assuming uniform distribution of soil N resources and soil microbial communities that mineralise organic N. However, organic N resources are largely concentrated in the topsoil (Balesdent *et al*. 2018), while mineral N, especially nitrate also occurs in deeper soil layers as a result of leaching (Lynch 2013). Similarly, microbial N mineralisation often occurs in distinct pockets of soil, so-called hotspots that are rich in organic N and provide favourable conditions for microbial activity (Hesselsøe *et al*. 2001; Hill *et al*. 2019). Localised root growth and exudation in response to spatial differences in nutrient availability has been reported in a range of plant species (in ‘t Zandt *et al*. 2015; Lemming *et al*. 2016; Paterson *et al*. 2006; Peñaloza *et al*. 2002). Hence, the maximisation of aboveground plant biomass requires spatial regulation of root growth and exudation to co-localise C investment with different soil N resources. Root architecture and the spatial distribution of soil N are explicitly represented in structural-functional root models such as *OpenSimRoot* and *CRootBox* (Postma *et al*. 2017; Schnepf *et al*. 2018). Combining these models with the dynamic regulation of root C partitioning presented here will allow to decipher the contribution of spatially confined root growth and exudation for maximizing aboveground plant biomass.

Acknowledging that plants adjust their belowground physiology to changes in the availability of different soil N resources is imperative to develop mechanistic understanding of plant responses to future soil environments. The model we present here is based on the central assumption that plants dynamically regulate C partitioning between root growth and exudation to optimise their N foraging behaviour under varying soil N availability. The explicit representation of belowground plant acclimations enables us to answer the question: “*What must a particular plant do to maximize its performance under given soil conditions?*”. Thus, our eco-evolutionary approach to modelling N foraging represents a paradigmatic shift away from models that rely on fixed plant physiological behaviour and ultimately ask: “*How does a particular plant perform under given soil conditions?*”. This paradigmatic shift is indispensable to generate hypotheses on optimal N foraging strategies in future soil environments characterised by more erratic N availability. Thereby, our N foraging model can guide the design of empirical studies on the effects of changing soil N resources on C and N fluxes in plant-soil systems governing plant growth and ecosystem productivity.

## ACKNOWLEDGEMENTS

We thank Stefano Manzoni (Stockholm University, Sweden) for feedback to an early version of the manuscript. TC acknowledges funding from the University of Nottingham (Nottingham Research Fellowship) and SJM is funded by BBSRC Project Designing Sustainable Wheat (BB/X018806/1) and (BB/X011003/1) and BBSRC BreakThru (BB/W008874/1). A small portion of this project was administered through EMSL, the Environmental Molecular Sciences Laboratory. EMSL is a Dept of Energy, Office of Science National User Facility.

## AUTHOR CONTRIBUTION

AC and TC conceived the study and developed the modelling framework with inputs from SJM. AC implemented the model, wrote the model code, and preformed all simulations. AC and TC wrote the original draft of the manuscript, SJM contributed to review and editing. Funding was acquired by TC.

## CODE AVAILABILITY

The MATLAB scripts used to perform simulations and to generate the presented figures can be accessed from https://doi.org/10.5281/zenodo.15367434.

