## Supplementary information for "Modelling belowground plant acclimation to low soil nitrogen – An eco-evolutionary approach"

**Supplemental Materials**

Supplemental Tables: 5

Supplemental Table S1: List of state variables and their mass balances equations

| C Mass balance | | | |
|---|---|---|---|
| **State variable** | **Unit** | **Equation** | |
| Leaf C | $gC\ m^{-2}$ | $\frac{dC_L}{dt} = A - R_L - S$ | (1) |
| Root C | $gC\ m^{-2}$ | $\frac{dC_R}{dt} = S - E - R_{R,g} - R_{R,m} - T_R$ | (2) |
| Exudate C | $gC\ m^{-2}$ | $\frac{dC_E}{dt} = E - U_E$ | (3) |
| SOM C | $gC\ m^{-2}$ | $\frac{dC_{SOM}}{dt} = -U_{SOM} + T_M + T_R$ | (4) |
| Microbial C | $gC\ m^{-2}$ | $\frac{dC_M}{dt} = U_E + U_{SOM} - R_M - T_M$ | (5) |

| N Mass balance | | | |
|---|---|---|---|
| **State variable** | **Unit** | **Equation** | |
| Leaf N | $gN\ m^{-2}$ | $\frac{dN_L}{dt} = p_N = \frac{A - R_L - S}{CN_L}$ | (6) |
| Root N | $gN\ m^{-2}$ | $\frac{dN_R}{dt} = U_N - \frac{E}{CN_E} - \frac{T_R}{CN_R} - p_N$ | (7) |
| Exudate N | $gN\ m^{-2}$ | $\frac{dN_E}{dt} = \frac{E}{CN_E} - \frac{U_E}{CN_E} = \frac{1}{CN_E}\frac{dC_E}{dt}$ | (8) |
| SOM N | $gN\ m^{-2}$ | $\frac{dN_{SOM}}{dt} = -\frac{U_{SOM}}{CN_{SOM}} + \frac{T_M}{CN_M} + \frac{T_R}{CN_R}$ | (9) |
| Microbial N | $gN\ m^{-2}$ | $\frac{dN_M}{dt} = \frac{U_E}{CN_E} + \frac{U_{SOM}}{CN_{SOM}} - \frac{T_M}{CN_M} - \phi_N$ | (10) |
| Mineral N | $gN\ m^{-2}$ | $\frac{dN_{min}}{dt} = \phi_N - U_N$ | (11) |

Abbreviation: SOM = Soil organic matter

Supplemental Table S2: Kinetic relations between C and N stocks and rates

| Symbol | Description | Expression |
|---|---|---|
| $A$ | Gross carbon assimilation rate of leaf | $A = N_L\left(1 - \frac{N_L}{K_{NL}}\right)(a - bC_P)$ |
| $A_{net}$ | Net photosynthesis rate | $A_{net} = A - R_L = \mathrm{CUE_L}A$ |
| $CUE_L$ | Carbon use efficiency of leaf | $CUE_L = \frac{A - R_L}{A} = \frac{G_L + S}{A}$ |
| $CUE_M$ | Carbon use efficiency of soil microorganisms | $CUE_M = \frac{U_E + U_O - R_M}{U_E + U_O} = \frac{G_M}{U_E + U_O}$ |
| $CUE_R$ | Carbon use efficiency of root | $CUE_R = \frac{S - R_{R,g}}{S} = \frac{G_R + E + R_{R,m}}{S}$ |
| $E$ | Root exudation rate | *Estimated* |
| $E_{max}$ | Maximum root exudation rate | $E_{max} = V_{max,R}^{E} * C_R$ |
| $G_L$ | Net carbon assimilation rate of leaf | $G_L = CUE_L A - S$ |
| $G_M$ | Net carbon assimilation rate of soil microorganisms | $G_M = CUE_M(U_E + U_{SOM})$ |
| $G_R$ | Net carbon assimilation rate of root | $G_R = CUE_R S - E - R_{R,m}$ |
| $I_N$ | Mineral N supply rate to microbial pool | $I_N = \frac{V_{max,M}^{N_{min}} N_{min}}{K_M^{N_{min}} + N_{min}}$ |
| $p_N$ | Leaf N uptake rate | $p_N = \frac{A - R_L - S}{CN_L}$ |
| $R_L$ | Leaf respiration rate | $R_L = (1 - CUE_L)A$ |
| $R_M$ | Microbial respiration rate | $R_M = (1 - CUE_M)(U_E + U_O)$ |
| $R_{R,m}$ | Root maintenance respiration rate | $R_{R,m} = r_M C_R$ |
| $R_{R,g}$ | Root growth respiration rate | $R_{R,g} = (1 - CUE_R)S$ |
| $S$ | Root C supply rate | *Estimated* |
| $T_M$ | Microbial necromass turnover rate | $T_M = \tau_M C_M$ |
| $T_R$ | Root turnover rate | $T_R = \tau_R C_R$ |
| $U_E$ | Exudate uptake rate by soil microorganisms | $U_E = \frac{V_{max,M}^{E} C_E C_M}{K_E + C_M + C_E}$ |
| $U_{N,pot}$ | Potential mineral N uptake rate by roots | $U_{N,pot} = \frac{V_{max,R}^{N_{min}} N_R N_{min}}{K_R^{N_{min}} + N_{min}}$ |
| $U_{SOM}$ | SOM uptake rate | $U_{SOM} = \frac{V_{max}^{SOM} C_{SOM} C_M}{K_{SOM} + C_M + C_{SOM}}$ |
| $\phi_N$ | N imbalance rate | $\phi_N = \frac{U_{SOM}}{CN_{SOM}} + \frac{U_E}{CN_E} - \frac{CUE_M(U_E + U_{SOM})}{CN_M}$ |

Supplemental Table S3: Mathematical representation of nitrogen foraging model as an optimal control problem

$$\text{Maximize}\, J(\boldsymbol{x}(\boldsymbol{t}), \boldsymbol{u}(\boldsymbol{t})) = \int_0^T G_L\, dt$$

Subject to:

$$\frac{\mathrm{d}\boldsymbol{x}}{\mathrm{d}t} = f(\boldsymbol{x}(t), \boldsymbol{u}(t), p, t)$$

$$\boldsymbol{x}(t_0, p) = \boldsymbol{x_0}$$

$$\boldsymbol{x}(t), \boldsymbol{u}(t) \geq 0$$

$$\boldsymbol{u}(\boldsymbol{t}) = [S(\boldsymbol{t}), E(\boldsymbol{t})]$$

$$G_R(t), G_L(t), S(t) \geq 0$$

$$0.2 \leq \frac{G_R(t)}{S(\boldsymbol{t})} \leq 1$$

$$0.2 \leq \frac{G_L(t)}{A_{net}(\boldsymbol{t})} \leq 1 \qquad (12)$$

$$0 \leq \frac{U_N(t)}{U_{N,pot}(\boldsymbol{t})} \leq 1$$

$$0.1 \leq \frac{E(t)}{E_{max}(\boldsymbol{t})} \leq 1$$

where, $J$ is the objective function, which is cumulative aboveground plant growth, $x$ is the state variable vector, $u$ is the vector of control variables being optimized (i.e., root C supply, $S(\boldsymbol{t})$, and exudation rates, $E(\boldsymbol{t})$), $p$ is model parameter vector. Lower bounds to rates were applied to avoid unrealistic predictions or $S$ and $E$.

Supplemental Table S4: List of model parameters, their values, and units.

| Symbol | Description | Value | Unit | Source |
|---|---|---|---|---|
| $a$ | Maximum nitrogen productivity | 1.64 | $\mathrm{gC\ gN^{-1}d^{-1}}$ | *Assumed* |
| $b$ | Shading factor | 0.001 | – | Ågren 1985; Baskaran *et al.* 2017 |
| $BD$ | Soil bulk density | 1.2 | $\mathrm{g\ cm^{-3}}$ | Panagos *et al.* 2024 |
| $CUE_R$ | Root carbon use efficiency | 0.6 | – | Herrmann & Colombi 2019 |
| $CUE_L$ | Leaf carbon use efficiency | 0.7 | – | Hansen *et al.* 2009 |
| $CUE_M$ | Microbial carbon use efficiency | 0.3 | – | Blagodatskaya *et al.* 2014; Mo *et al.* 2021 |
| $CN_{SOM}$ | C-to-N ratio of SOM | 10 (8-25) | $\mathrm{gC\ gN^{-1}}$ | Tipping *et al.* 2016 |
| $CN_R$ | C-to-N ratio of roots | 50 | $\mathrm{gC\ gN^{-1}}$ | Zhang *et al.* 2020 |
| $CN_M$ | C-to-N ratio of soil microorganisms | 10 | $\mathrm{gC\ gN^{-1}}$ | Zhang & Elser 2017 |
| $CN_L$ | C-to-N ratio of leaves | 21 | $\mathrm{gC\ gN^{-1}}$ | Zhang *et al.* 2020 |
| $CN_E$ | C-to-N ratio of exudates | 10000 | $\mathrm{gC\ gN^{-1}}$ | *Assumed* |
| $K_E$ | Half-saturation constant for exudate uptake kinetics | 3.3 | $\mathrm{gC\ m^{-2}}$ | Approximated from Vinolas *et al.* 2001 |
| $K_M^{N_{min}}$ | Half-saturation constant for mineral N supply rate to microbial pool | 1 | $\mathrm{gN\ m^{-2}}$ | *Assumed* |
| $K_{NL}$ | plant N carrying capacity | 75 | $\mathrm{gN\ m^{-2}}$ | *Assumed* |
| $K_R^{N_{min}}$ | Half saturation constant for root mineral N uptake rate | 20 | $\mathrm{gN\ m^{-2}}$ | *Assumed* |
| $K_{SOM}$ | Half saturation constant for SOM uptake rate | 6443 | $\mathrm{gC\ m^{-2}}$ | Abramoff *et al.* 2022 |
| $Root\ C\%$ | C content in dry root biomass | 0.4 | $\mathrm{gC\ g^{-1}}$ dry weight root | Ma *et al.* 2018 |
| $V_{max,M}^{E}$ | Maximum exudate uptake rate by soil microorganisms | 5.3 | $\mathrm{d^{-1}}$ | Approximated from Vinolas *et al.* 2001 |
| $V_{max,R}^{E}$ | Maximum root exudation rate constant | 0.035 | $\mathrm{d^{-1}}$ | *Assumed* |
| $V_{max,M}^{N_{min}}$ | Max. mineral N uptake rate by soil microorganisms | 15 | $\mathrm{d^{-1}}$ | *Assumed* |
| $V_{max,R}^{N_{min}}$ | Maximum mineral N uptake rate by roots | 0.25 | $\mathrm{d^{-1}}$ | Griffiths & York 2020; York *et al.* 2016 |
| $V_{max}^{SOM}$ | Maximum SOM uptake rate | 0.15 | $\mathrm{d^{-1}}$ | *Assumed* |
| $\tau_M$ | Microbial turnover rate constant | 0.0137 | $\mathrm{d^{-1}}$ | *Assumed* |
| $\tau_R$ | Root turnover rate constant | 0.001 | $\mathrm{d^{-1}}$ | Gill & Jackson 2000 |

Abbreviation: SOM = Soil organic matter

Supplemental Table S5: Initial condition of state variables in base, low mineral N, and high SOM C:N ratio scenarios.

| **Initial condition (g C $m^{-2}$ or g N $m^{-2}$ or [-])** | **Base scenario** | **Low mineral N scenario** | **High SOM C:N ratio scenario** |
|---|---|---|---|
| Leaf C | 50 | 50 | 50 |
| Root C | 62.5 | 62.5 | 62.5 |
| Exudate C | 0.5 | 0.5 | 0.5 |
| SOM C | 7200 | 7200 | 7200 |
| Microbial C | 2.8 | 2.8 | 2.8 |
| Microbial N | 0.28 | 0.28 | 0.28 |
| Mineral N | 20 | 10 | 20 |
| SOM C:N ratio | 10 | 10 | 20 |

Abbreviation: SOM = Soil organic matter

**Cited references**


Abramoff, R.Z., Guenet, B., Zhang, H., Georgiou, K., Xu, X., Viscarra Rossel, R.A., *et al.* (2022). Improved global-scale predictions of soil carbon stocks with Millennial Version 2. *Soil Biol. Biochem.*, 164, 108466.

Ågren, G.I. (1985). Theory for growth of plants derived from the nitrogen productivity concept. *Physiol. Plant.*, 64, 17–28.

Baskaran, P., Hyvönen, R., Berglund, S.L., Clemmensen, K.E., Ågren, G.I., Lindahl, B.D., *et al.* (2017). Modelling the influence of ectomycorrhizal decomposition on plant nutrition and soil carbon sequestration in boreal forest ecosystems. *New Phytol.*, 213, 1452–1465.

Blagodatskaya, E., Blagodatsky, S., Anderson, T. & Kuzyakov, Y. (2014). Microbial Growth and Carbon Use Efficiency in the Rhizosphere and Root-Free Soil. *PLoS One*, 9, e93282.

Gill, R.A. & Jackson, R.B. (2000). Global patterns of root turnover for terrestrial ecosystems. *New Phytol.*, 147, 13–31.

Griffiths, M. & York, L.M. (2020). Targeting Root Ion Uptake Kinetics to Increase Plant Productivity and Nutrient Use Efficiency. *Plant Physiol.*, 182, 1854–1868.

Hansen, L.D., Thomas, N.R. & Arnholdt-Schmitt, B. (2009). Temperature responses of substrate carbon conversion efficiencies and growth rates of plant tissues. *Physiol. Plant.*, 137, 446–458.

Herrmann, A.M. & Colombi, T. (2019). Energy use efficiency of root growth – a theoretical bioenergetics framework. *Plant Signal. Behav.*, 14, 1685147.

Ma, S., He, F., Tian, D., Zou, D., Yan, Z., Yang, Y., *et al.* (2018). Variations and determinants of carbon content in plants: a global synthesis. *Biogeosciences*, 15, 693–702.

Mo, C., Jiang, Z., Chen, P., Cui, H. & Yang, J. (2021). Microbial metabolic efficiency functions as a mediator to regulate rhizosphere priming effects. *Sci. Total Environ.*, 759, 143488.

Panagos, P., De Rosa, D., Liakos, L., Labouyrie, M., Borrelli, P. & Ballabio, C. (2024). Soil bulk density assessment in Europe. *Agric. Ecosyst. Environ.*, 364, 108907.

Tipping, E., Somerville, C.J. & Luster, J. (2016). The C:N:P:S stoichiometry of soil organic matter. *Biogeochemistry*, 130, 117–131.

Vinolas, L.C., Healey, J.R. & Jones, D.L. (2001). Kinetics of soil microbial uptake of free amino acids. *Biol. Fertil. Soils*, 33, 67–74.

York, L.M., Silberbush, M. & Lynch, J.P. (2016). Spatiotemporal variation of nitrate uptake kinetics within the maize ( Zea mays L .) root system is associated with greater nitrate uptake and interactions with architectural phenes. *J. Exp. Bot.*, 67, 3763–3775.

Zhang, J. & Elser, J.J. (2017). Carbon:Nitrogen:Phosphorus Stoichiometry in Fungi: A Meta-Analysis. *Front. Microbiol.*, 8, 1–9.

Zhang, J., He, N., Liu, C., Xu, L., Chen, Z., Li, Y., *et al.* (2020). Variation and evolution of C:N ratio among different organs enable plants to adapt to N-limited environments. *Glob. Chang. Biol.*, 26, 2534–2543.
